# Optimising play for learning risky behaviour

**DOI:** 10.1101/2025.07.15.664889

**Authors:** Dharanish Rajendra, Chaitanya S. Gokhale

## Abstract

Animals adapt their behaviour to current environmental conditions to enhance survival and reproductive success. While longterm adaptation occurs through evolutionary processes acting on heritable variation, individuals can also adapt within their lifetime via learning. Learning is particularly advantageous in environments that are uncertain or fluctuate across a lifespan or a few generations. However, reliance on individual learning entails a critical risk. Juveniles may begin life poorly adapted to their surroundings, requiring exploration to learn. Such an approach can be costly and dangerous, especially for species engaging in risky activities such as hunting dangerous prey. We explore how early-life learning in a protected environment, such as one buffered by parental care, can facilitate effective behavioural adaptation in later, riskier contexts. As a representative case, we model the decision-making process of a predator hunting both safe and dangerous prey. We analyse decisionmaking dynamics through reinforcement learning, extending beyond classical dynamic programming approaches. Our results show that experiences in a juvenile’s early environment can generalise to a distinct adult environment, provided there is sufficient structural similarity between them. Our findings demonstrate that incorporating structured play or safe exploration in early life can significantly enhance the performance of learningbased adaptation in dangerous environments.

## 1. Introduction

Learning and evolution represent fundamental yet distinct mechanisms of biological adaptation (1, 2). While evolution shapes traits across generations through genetic selection (3), learning enables behavioural flexibility within an individual’s lifetime (4). This dual adaptive system is especially advantageous in variable environments, where learning can compensate for the relatively slow pace of evolutionary change (5, 6). Learning also confers benefits in complex environments by allowing individuals to tailor behaviour to local contingencies (7). However, both learning and the capacity for learning are costly (8–10). Organisms that rely heavily on learning produce individuals with the potential to acquire adaptive behaviours but who are initially maladapted to their environment (11). This creates an apparent paradox: individuals must survive a vulnerable, uninformed phase before they can learn how to behave adaptively.

This paradox is particularly acute in the context of risky behaviours, where mistakes can be lethal (8, 12). Predators that hunt dangerous prey exemplify such scenarios, a strategy observed across multiple animal taxa (13). Effective predation on dangerous prey typically requires precise and often complex behavioural repertoires, which are not innate but must be acquired through individual experience or social learning (14). The predators must also assess the situation — the dangerousness of a prey, the potential food benefit and also its own capture probability and energy levels – before making the decision of hunting. Jumping spiders exemplify this situation. They hunt a wide variety of dangerous prey, can identify and distinguish between them and learn through trial and error the strategies that work for different kinds of prey (15). They also take into account their own energy level before choosing prey to hunt (16, 17). Paradoxically, theory suggests that hunting dangerous prey can emerge as an evolutionarily stable strategy even when safer alternatives exist (18, 19). Yet the mechanisms through which naive individuals acquire these high-stakes behaviours remain poorly understood.

Extended parental care, particularly prevalent in species with advanced learning capabilities (20, 21), may resolve this paradox. Developmental stages such as infancy and juvenile phases may provide critical windows for safe learning (22). The phases can be further supported by parental behaviours that buffer young from environmental hazards (23). However, this gives rise to a key question: to what extent can behaviours learned in protected environments generalise to real-world, high-risk contexts?

Some previous studies have shown that if the costs of learning are outweighed by the benefits of adapting to a changing environment through individual or social learning, then it is likely to be advantageous and evolve (24, 25). However, these models assume a very simple form of behaviour and learning and do not include the multi-step decision process which animals have to go through. The learning process in these models is abstract and assumes that a sufficient proportion of juveniles somehow learn the correct behaviour without dying. However, this is very unlikely when the individuals have to learn a multi-step state-based policy. Here, we explore one of the ways that the cost can be overcome through a modification of the juvenile environment by its parents.

In this study, we investigate how early-life learning in protected settings influences the development of prey choice strategies in predators. Using a combination of dynamic programming (26, 27) and reinforcement learning frameworks (28), we model the acquisition of hunting strategies targeting both safe and dangerous prey. Dynamic programming has been successfully applied not only to prey choice & foraging (18) and other animal behaviours (29), but also to a range of other ecological and evolutionary contexts (30), including the evolution of mimicry and signal learning (31). Similar reinforcement learning methods have been applied previously to model animal behaviour and learning, notably by McNamara and Leimar (32, 33). Animals may make decisions and perform actions while considering long-term impacts in addition to short-term benefits. Crucially, reinforcement learning captures how animals make decisions by accounting for both immediate and long-term consequences. That is, current actions influence future states and rewards, something often ignored in theoretical models of learning or its evolution (but see (34–36) for exceptions). For a broad comparison between reinforcement learning and the more classical dynamic programming methods, see (37).

Specifically, we explore (i) the conditions under which behavioural skills learned in protected environments can transfer to more dangerous ones, (ii) how environmental danger levels shape learning efficiency and performance in adulthood, and (iii) the relation between the properties of the external and protected environments which result in a good transfer of skills and performance.

We find that learning in protected environments can significantly improve an individual’s ability to adapt to later risky environments, especially when there is structural similarity between the two. For some environmental conditions, simply providing a safe, protected environment in the juvenile phase is sufficient to promote learning. For other environmental conditions, a more involved strategy of providing a highly protected environment for the initial part of the juvenile phase and then switching to a less protected environment increases the performance and adaptation of the individual in adulthood. There exists an optimal duration of juvenile learning that maximises adult reproductive success, balancing the trade-off between early safety and delayed experience.

We begin by describing the behavioural situation we are investigating with a baseline of expected optimal behaviour. That is followed by a description of the modelling framework of reinforcement learning. We then present the results on the performance of an individual given the juvenile environments it experiences. An important quantity is the time it takes to learn in the adult state the adult re-learning time. Finally, we discuss the implications of our findings for behavioural development, life-history evolution, and the evolution of extended parental care and of complex learning mechanisms.

## 2 Decision-making and energy dynamics

We investigate prey choice dynamics through a stochastic energy budget model of a predator encountering heterogeneous prey types (38). Specifically, we model a predator’s foraging strategy confronting two distinct prey categories: safe and dangerous. The predator’s energetic state is conceptualized as a jump process (39), characterized by discrete transitions in metabolic potential (Fig. 1). Thus, the predator’s energy remains stationary over discrete time intervals, punctuated by probabilistic transitions that reflect metabolic fluctuations, hunting outcomes, and physiological variability. These transitions represent changes in energetic state induced by the predator’s decisions and capture the differential risks of pursuing safe versus dangerous prey (40). The jump process framework allows us to model the complex, non-linear dynamics of predator energy acquisition under uncertainty (41) (detailed in App. A.1).

**Fig. 1.**
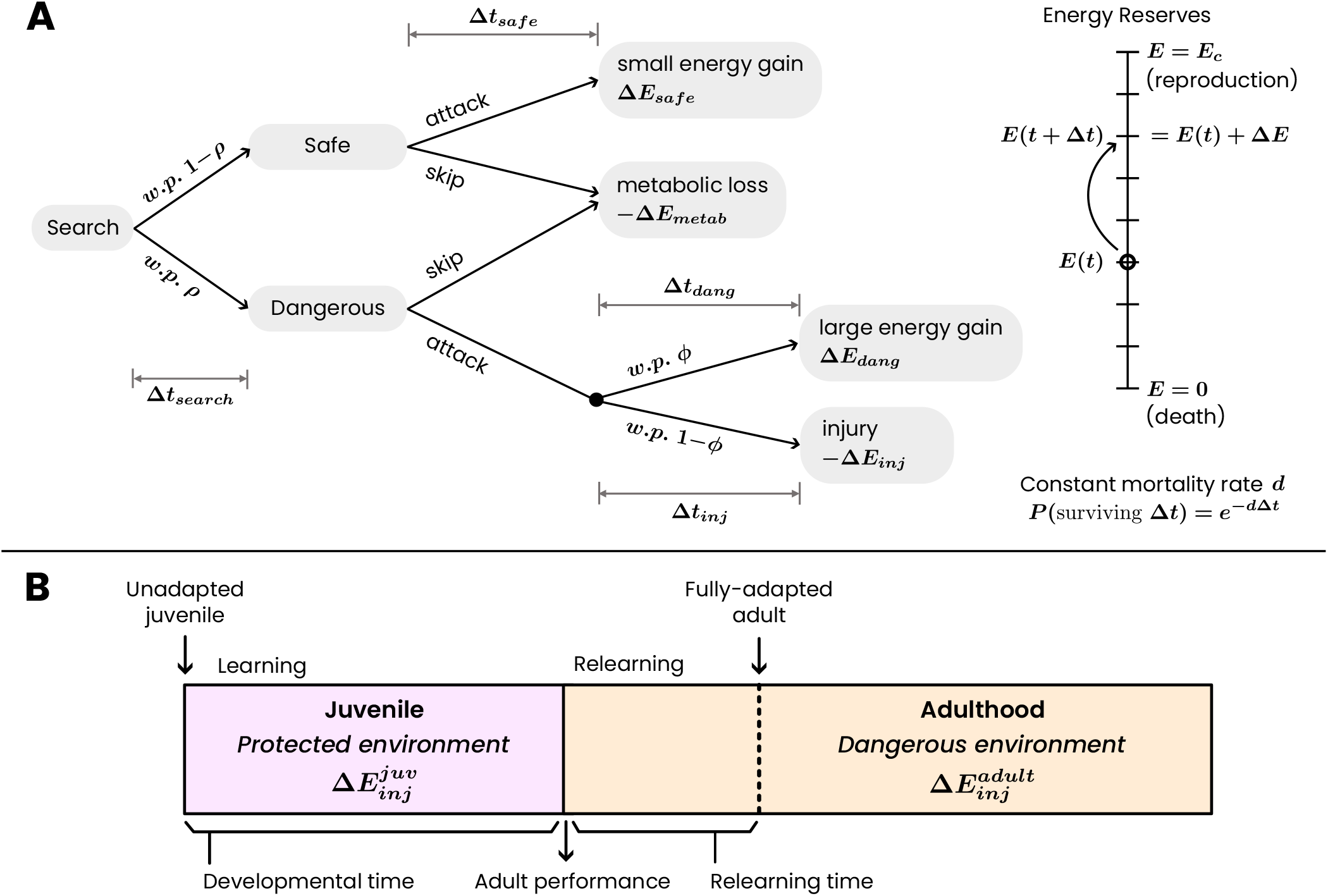
Depiction of the model processes. **A**. Flow chart depicting the stochastic decision-making model of predator foraging behaviour and energy dynamics. At each step of the process, the predator begins with the searching step. It encounters safe or dangerous prey with a probability according to the availability of dangerous prey *ρ*, and after a searching time depending on the total prey abundance. Then the predator can choose to skip or attack the encountered prey. The decision results in energy gains or losses (Δ*E*_*i*_) for action *i* with time investments (Δ*t*_*i*_), and potential injury risks. The predator’s energy reserves (*E*(*t*)) changes stochastically, with a constant mortality rate (*d*) influencing survival probability. The predator succeeds probabilistically against a dangerous prey with a probability depending on its capture probability *ϕ*. Upon failure, it loses energy Δ*E*_*inj*_. **B**. Illustration of the learning processes that an individual experiences in their lifetime. A juvenile grows up in a protected environment created by its parents, in which it learns to adapt to and perform well. As an adult, it is exposed to the truly dangerous elements of the environment, where it must relearn and re-adapt.

The animal’s energy level varies between zero and a fixed maximum value, denoted as *E*_*c*_ (Fig. 1A). The goal of the animal is to reach the maximum energy level, which corresponds to a fitness benefit, and to avoid the lower energy limit. Each time step consists of prey search, encounter, decision, and outcome, with consequences for energy level and elapsed time. The predator searches for prey and encounters an individual after a delay Δ*t*_*search*_, contingent upon the availability of prey in the environment. Upon encountering prey, which could be dangerous with probability *ρ* and safe with probability 1−*ρ*, the predator assesses its danger level and makes a decision. The predator has the option to either attack the prey or abstain. Based on its decision, the predator’s energy level undergoes a stochastic change, and a time interval elapses before it resumes the search process. This process repeats until the energy level either reaches the reproduction boundary or reaches zero.

If the predator opts to skip the prey it has encountered, it neither gains nor loses any energy. However, it has already expended energy on searching. Consequently, it incurs a metabolic cost of Δ*E*_*metab*_, which depends on its metabolic rate and the duration of the search. This energy reduction occurs after the search time.

If the prey is safe and the predator attacks, the predator handles it over a time Δ*t*_*safe*_ and gains a small amount of energy Δ*E*_*safe*_. The net energy gain is the difference between the prey’s value and the energetic cost of search and handling. The process transitions to the next step after time Δ*t*_*search*_ +Δ*t*_*safe*_.

If the prey is dangerous and attacked, the outcome is probabilistic. The attack succeeds with a probability *ϕ* depending on the prey’s capture probability. On a successful attack, the net energy gained, Δ*E*_*dang*_, is substantial (generally larger than the safe prey) after a handling time Δ*t*_*dang*_. If the attack fails (with probability 1− *ϕ*), we interpret it to be due to the prey retaliating. The predator is then injured and must expend energy Δ*E*_*inj*_ and time Δ*t*_*inj*_ to recover. It is possible that the injury is so grave that it is not possible to recover from it, and the predator dies. This is captured by the case when the recovery energy is greater than the current energy level of the predator, i.e. Δ*E*_*inj*_ *> E*(*t*).

All energies and times are, in general, random variables with their own distributions, means, and variances. Of importance are the inequalities between these variables between the safe and dangerous prey. Moreover, the predator is subject to a constant background mortality rate *d*, which may cause death regardless of energy level. Consequently, the survival probability depends on the duration of time between events. Therefore, the survival of an individual from one step to the next is probabilistically determined by the mortality rate and the time interval between these steps. Mathematically, the probability of surviving an interval of time Δ*t* is *s*(Δ*t*) = exp(*−d*Δ*t*).

To simplify analysis, we will not directly consider the predator’s individual actions of attacking and skipping the two prey types, but rather consider three compound behaviours that the predator can choose between and execute at each decision-making step:

1. Indiscriminate attack: attack the prey encountered regardless of its type.
2. Risk-prone attack: attack the prey only if it is of the dangerous type
3. Risk-averse attack: attack the prey encountered only if it is safe.

We must reiterate that these are not identities of the agent but behaviours that it can choose from at each step. That is, for example, a predator can adopt a strategy which is to be indiscriminate at low energy levels and risk-prone at higher energy levels. Such an energy-dependent (in general, statedependent) strategy is known as a *policy*. A policy *π* is formally defined as an energy-dependent rule: it specifies the action *A* = *π*(*E*) that the predator should take when at energy level *E*. This is equivalent to a behavioural strategy that adapts to the current energetic state, and can vary across the predator’s lifespan.

The predator reproduces when *E* = *E*_*c*_. This reward received by the predator makes this state directly valuable. All other energy states acquire an indirect value through their probability of reaching *E*_*c*_ before death. Such a value is quantified by the *value function V* (*E*), which is defined as the expected reproductive payoff when starting from energy level *E*.

An optimal policy, *π*\*, maximises the value function *V* (*E*), and hence the expected reproductive payoff, for each energy level (see App. A.2). This optimal policy can be computed using the value iteration method of dynamic programming (26, 27), which is well-suited to solving such sequential decision problems.

### 2.1. Optimal foraging policies across risk profiles

We compute the optimal foraging policy over a range of environmental and individual parameters, specifically dangerous prey availability (*ρ*) and predator capture probability (*ϕ*) (Fig. 2). The policy is a vector of decisions across energy levels from 1 to *E*_*c*_ (we use *E*_*c*_ = 100), where each entry specifies the best action at that energy state. For visualisation purposes, we reduce this policy vector into a tuple: (*in, rp, ra*), denoting the number of energy levels where the predator should attack indiscriminately, be risk-prone, or be risk-averse, respectively. This tuple is then mapped to a three-component colour and displayed as a heatmap in Fig. 2.

**Fig. 2.**
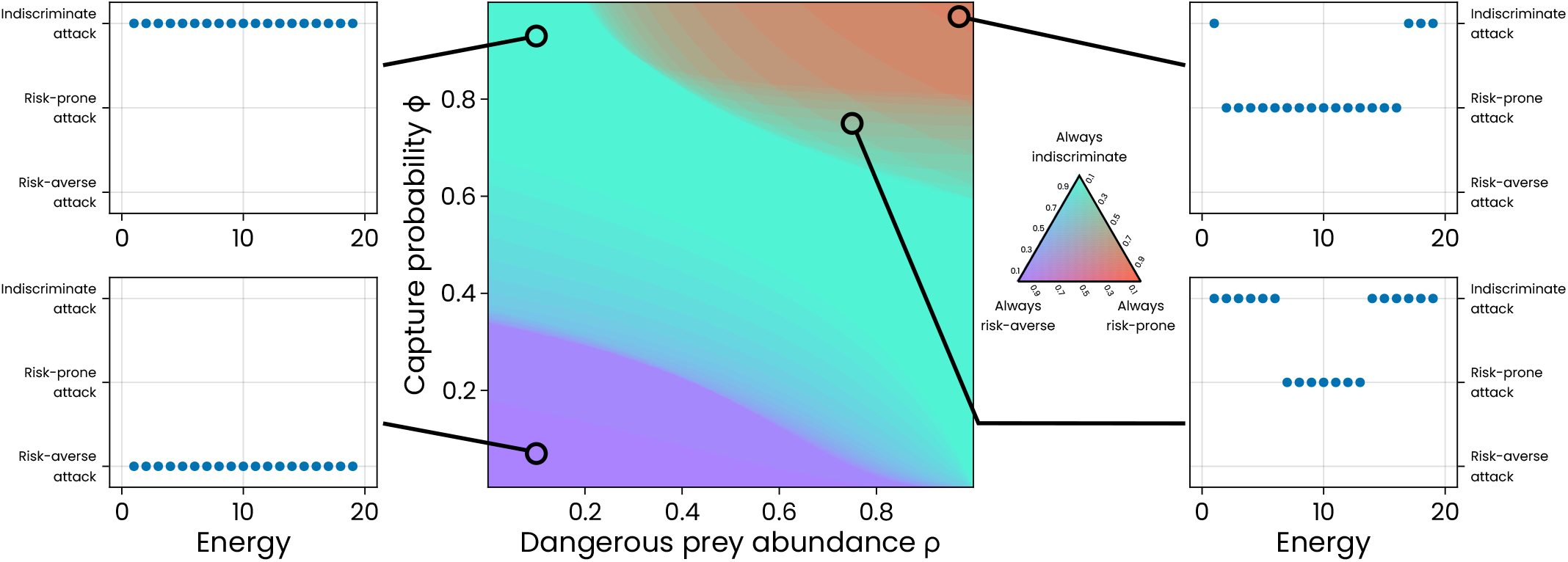
Optimal policy. A representation of the optimal policy for a range of parameter values and example policies for different regions. The colour of the points in the heatmap depicts the composition of the three compound behaviours of the policy.

The resulting phase diagram reveals three broad behavioural regimes. In the bottom-left region of the heatmap, corresponding to low capture probability *ϕ* and intermediate dangerous prey availability *ρ*, the optimal strategy is to exclusively specialise on safe prey across all energy levels. This reflects the high risk and low success rate of attacking dangerous prey under these conditions. In contrast, the top-right region, high *ϕ* and high *ρ*, favours a strategy of specialising on dangerous prey for most energy levels. The predator’s high skill and the abundance of high-energy prey make it optimal to fully rely on the risky but rewarding option. This is true, however, only for very high capture probabilities. For slightly lower capture probability and abundance of dangerous prey, it is a mixture of specialising on dangerous prey and generalising. In the top-left and bottom-right corners, where one of the quantities is low and the other is high, it becomes optimal to generalise regardless of energy level. In general, between these extremes lies a region of mixed policies. Here, the policy is not homogeneous: some energy levels favour generalising, while others favour specialising on a single prey type. Interestingly, across all parameter combinations, we observe no optimal policies that include both specialising on dangerous and safe prey. This suggests that mixed specialisation across prey types is never favoured. We also find regions dominated by generalist strategies, where the predator always attacks indiscriminately, regardless of the prey type.

This pattern can be understood by considering the trade-off between starvation risk and reproductive gain. When the capture probability (*ϕ*) and availability of dangerous prey (*ρ*) are both low, the costs of attacking dangerous prey are greater than the potential benefits, and it is optimal to ignore them entirely. As the capture probability increases from here, incorporating dangerous prey at intermediate energy levels is beneficial, as it would provide a chance to rapidly increase energy without a major chance of starvation. However, attacking a dangerous prey at low energy levels is still not optimal, as it can result in death from injury. As the capture probability increases further, including dangerous prey in the diet at all energy levels is beneficial, as the risk of fatal injuries at low energy levels has reduced. At high hunting abilities, beyond a certain threshold of the dangerous prey abundance, it becomes beneficial to ignore safe prey altogether. At this stage, skipping safe prey and waiting until a dangerous prey is encountered provides a higher rate of energy gain. This then transitions to complete dependence on dangerous prey for very high capture probability and dangerous prey availability. When the availability of dangerous prey is high, but the capture probability is low, it makes sense for the predator to attack any prey it encounters, as waiting for safe prey may result in it not getting any food at all and starving.

These are the optimal behaviours that an individual must perform in different environmental conditions in order to maximise its reproductive payoff. We assume that these prey choice behaviours are not genetically determined but learnt through trial and error throughout an individual’s life. In the next section, we detail how such a learning process is modelled and then move on to investigate how juvenile play can benefit their learning.

## 3. Reinforcement Learning

We use Temporal Difference (TD) reinforcement learning to model how a predator learns about its environment and develops an optimal foraging policy (28). TD learning is a class of algorithms that updates estimates of future reward by comparing successive predictions and capturing how the difference in value between states informs learning. This approach has been shown to reflect underlying neural mechanisms of animal learning (42, 43).

In our system, the learning agent estimates the expected value of each action at each energy level. These estimates are updated based on actual experiences during interaction with the environment. Specifically, updates occur when outcomes deviate from expectations: if a chosen action yields a higher reward than previously estimated, the agent increases the corresponding action-value proportionally to this discrepancy. The magnitude of this change is governed by the learning rate *α*. This can also be thought of as the rate at which the individual absorbs new information. The agent’s behaviour is guided by an *ϵ*-greedy policy: most of the time, it selects the action with the highest estimated value at its current energy level, but with a small probability *ϵ*, it explores other actions at random. Such a policy allows for exploring different (previously unknown) possibilities. As learning proceeds, the policy stochastically approaches the optimal one. Provided the learning rate is sufficiently small or gradually decays, convergence to the optimal policy is guaranteed in the long run.

At the start of an *experiment*, the agent is initialised with a random policy and a random energy level. The learning process continues until it reaches one of the boundaries (*E* = 0 or *E* = *E*_*c*_) and an *episode* is completed. Then, a new episode starts with a random initial energy, but the individual retains the learning from the previous round. Each learning run consists of multiple episodes. The process continues until a fixed number of steps/episodes are completed, or it reaches a learning target. At this point, one experimental run is considered to be completed. Multiple (10,000) independent experimental runs are performed for each parameter value in order to obtain the mean and variance for the metrics of interest.

In a constant environment, the agent eventually learns the deterministic optimal policy and value function (see Fig. A.1A). However, we are interested in the time to reach the optimal policy. This learning time (measured as the number of steps) depends on both the environmental parameters (Fig. A.1B) and the learning algorithm’s hyperparameters, and varies across realisations due to the stochastic nature of the learning process (the details of the learning algorithm are outlined in App. A.3).

### 3.1. Play in a protected environment

Many species provide parental care that creates a relatively protected environment during early development. In such contexts, juveniles can engage in learning and behavioural exploration with reduced risk. The goal of learning during this juvenile phase is to become well-adapted to the conditions provided by the parent. However, once parental protection ends, the juvenile transitions into a potentially hostile adult environment, where previously learned behaviours may no longer suffice. In this scenario, the individual must adapt once more and unlearn and/or relearn now through experience in a dangerous and unprotected environment. This is the scenario that we capture in the model.

Our model captures this developmental transition by dividing the individual’s life into two distinct stages: a juvenile phase and an adult phase. The environment experienced by the animal is constant within each phase but can be potentially different between phases. The key feature of this setup is that the juvenile phase provides an opportunity for learning in a safer and more protected context, while the adult phase presents the true conditions under which survival and reproduction depend. We vary various quantities of the juvenile phase and environment, such as its duration, dangerousness (quantified by the cost of injury Δ*E*_*inj*_), prey abundance *ρ*, and capture probability *ϕ* of dangerous prey. These parameters allow us to explore how different qualities of the early environment affect subsequent adaptation. To assess learning performance, we measure the time the animal requires to adapt to the adult environment by reaching the optimal policy (within a specified error threshold). We term this as the relearning time. This learning process is depicted in Fig. 1B

### 3.2. Adult performance

Once the individual transitions from the juvenile to the adult phase, their performance in the adult environment becomes critical for reproductive success. We quantify performance as the average reward obtained per episode in the adult environment immediately after the juvenile phase. This value is then normalised to lie between two benchmarks: the performance of a randomly selected policy and that of the optimal policy in the adult environment.

In general, individuals who spend more time learning during the juvenile phase achieve higher adult performance. However, this outcome strongly depends on the match or mismatch between the juvenile and adult environments (see Fig. 3). When the juvenile environment is identical to the adult environment (i.e., both are dangerous), performance improves steadily with developmental time and can approach optimality. In contrast, when the juvenile environment is a more protected proxy for the adult environment, the developmental trajectory of performance can differ substantially.

**Fig. 3.**
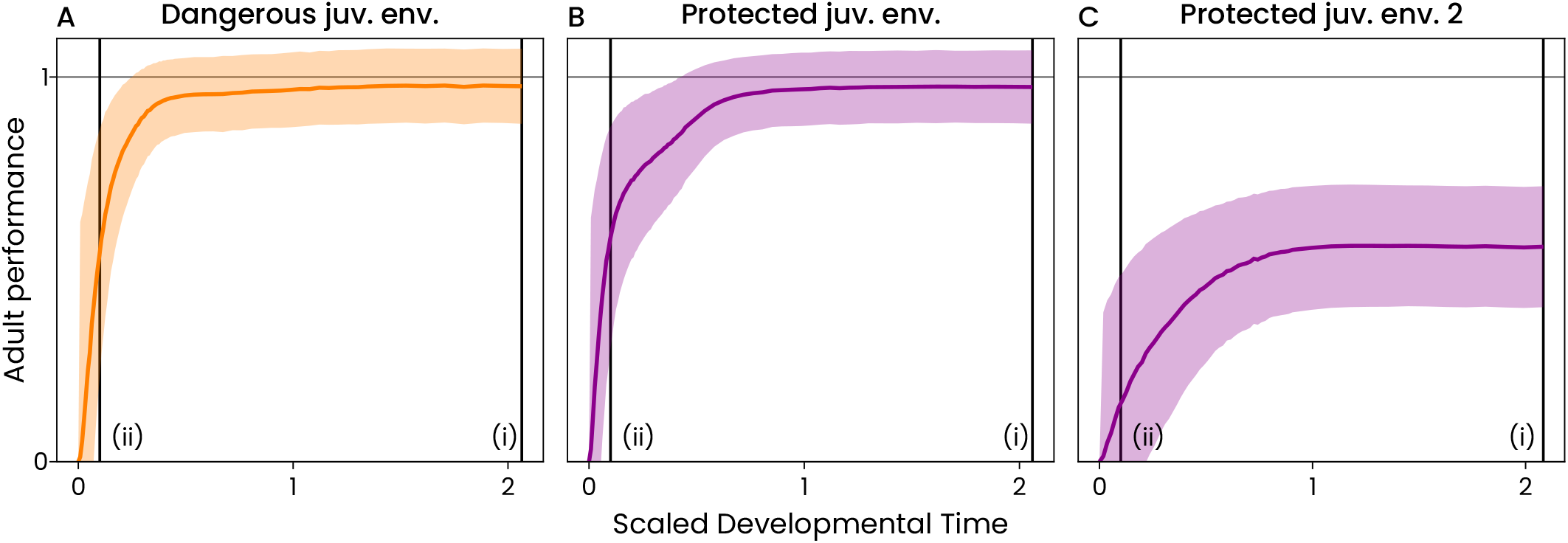
Adult performance is plotted against the developmental time the duration of the juvenile phase. The developmental time is scaled by the learning time in the dangerous environment without a juvenile phase. **A**. The juvenile and adult environments are the same environment dangerous (*ρ* = 0.9, *ϕ* = 0.6, Δ*E*_*inj*_ = 4). The adult performance increases with developmental time and saturates close to the maximum value. **B**. The juvenile environment is protected, but the adult environment is dangerous. Hence, as a juvenile, there is no injury cost upon failing to hunt a dangerous prey 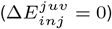. In this case, the adult performance increases with developmental time, but at a slower pace. However, it does approach the optimum. **C**. The juvenile environment in this case is protected, without any injury cost of failure 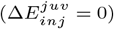. The adult environment is also dangerous but different from the two left panels (same Δ*E*_*inj*_ = 4 as the other panels but different *ρ* = 0.5 and *ϕ* = 0.2). In this case, the adult performance saturates well below the optimal performance. The bands in all panels show the standard deviation. Vertical lines are shown for (i) the longest developmental time (*>* 2) and (ii) a developmental time of 0.1. The adult performance is compared across dangerous and protected juvenile environments for these time points in Fig. 4A, B.

The adult performance for long developmental times may or may not be close to the optimal performance for the adult environment. Whether adult performance after extended development reaches the optimal level depends on several factors such as the abundance of dangerous prey (*ρ*), capture probability (*ϕ*), and the protection level of the juvenile environment (proxied by the injury cost Δ*E*_*inj*_). As shown in Fig. 4A, there is a region in the parameter space in which the maximum performance from the protected juvenile environment is significantly worse than the maximum performance from the dangerous juvenile environment. This discrepancy increases as the juvenile environment becomes more protected (i.e., as *E*_*inj*_ becomes smaller). Nevertheless, this outcome is not universal. There remains a substantial region of parameter space in which protected juvenile environments can support learning trajectories that eventually reach performance levels comparable to those developed in dangerous environments. This observation highlights that not all mismatches lead to maladaptive outcomes, particularly when adult challenges are not too dissimilar from juvenile experiences.

**Fig. 4.**
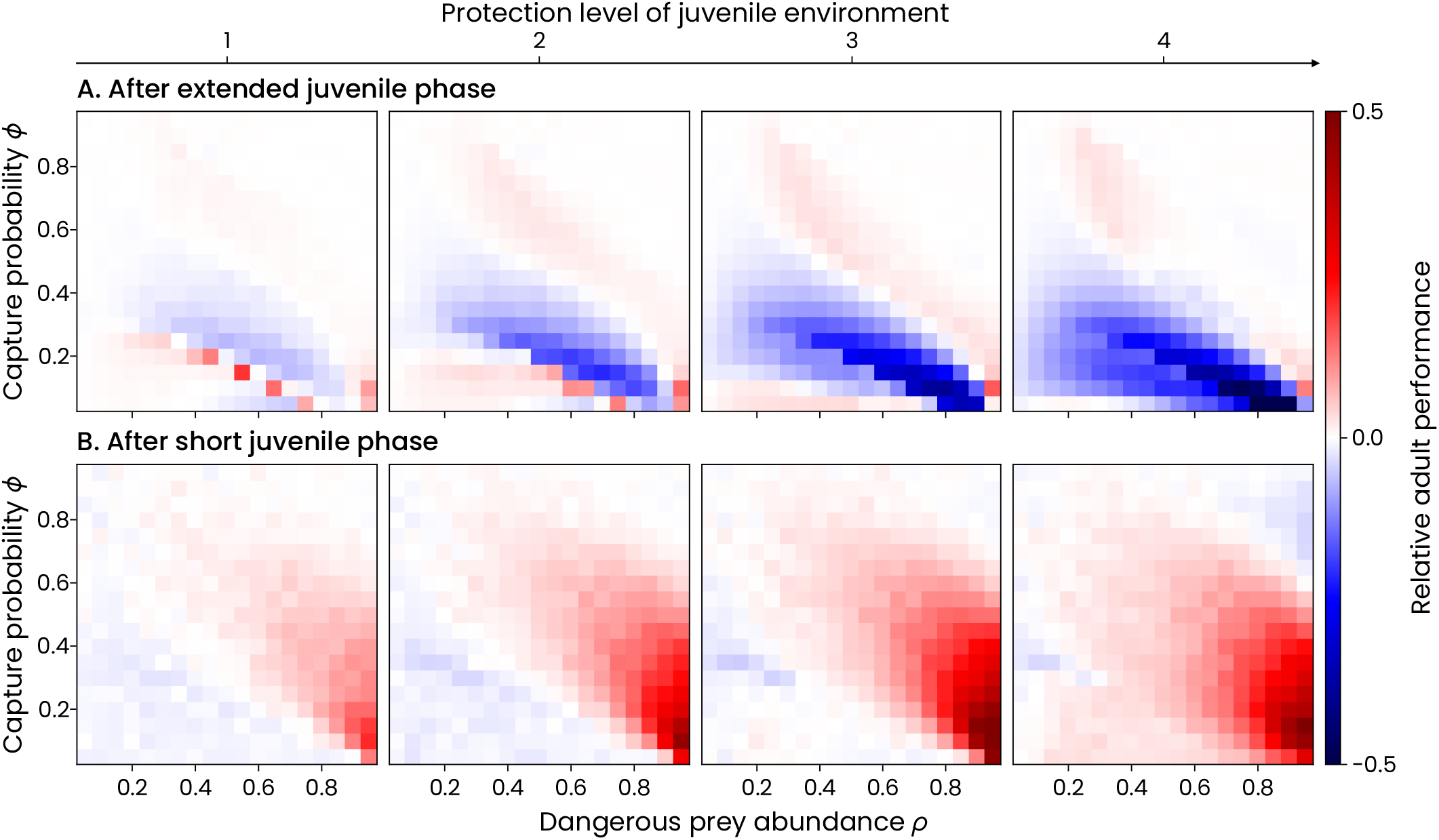
Adult performance after different developmental times. Adult performance is shown as a function of dangerous prey abundance (*ρ*), capture probability (*ϕ*), and juvenile environment protection level, measured by the reduction in injury cost from the dangerous to the protected environment 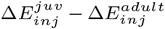. The corresponding adult environment for each panel has the danger level of 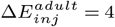. **A**. Adult performance is measured after an extended juvenile phase, i.e. a long developmental time. In the upper right triangle, the adult performance reaches close to the optimal performance (white/clear regions) for all levels of protection. In other areas, adult performance falls short of optimal and becomes progressively worse as the juvenile environment becomes more protected. **B**. Relative adult performance after a short juvenile phase with a developmental time of 0.1. Positive values (red) indicate that a short juvenile phase in the protected environment leads to a higher adult performance compared to a similar juvenile phase in the dangerous environment. Negative values (blue) indicate the opposite. White regions indicate similar adult performance between protected and unprotected juvenile environments. There is a sizeable region of parameter space where a protected environment leads to a higher adult performance for short juvenile durations. This region expands and becomes more prominent as the protection level of the juvenile environment increases.

Nevertheless, the difference between adult performance from a protected juvenile environment and from a dangerous juvenile environment is not the same over developmental time (see App. A.4). For adults coming from a protected environment, the performance can increase more rapidly than those from a dangerous juvenile environment for small developmental times, but saturate at a lower level. This is shown in Fig. A.2, where the relative adult performance is plotted for a range of developmental times and for a range of environmental parameter values. For low developmental times (see Fig. 4B), there is a large region of parameter space for all levels of protection of the juvenile environment, for which the adult performance is better than that from the dangerous juvenile environment. The size of this region is larger for more protected juvenile environments and decreases as the developmental time increases. As developmental time increases, a different region becomes more prominent, where the adult performance following a protected juvenile phase is worse than that following an unprotected juvenile phase. And even for long developmental times, this region still exists. This means that for these parameter values, the adult performance saturates significantly below the optimal performance, like in Fig. 3C.

### 3.3. Adult re-learning time

The second metric we use to quantify learning is re-learning time in the adult phase, where the individual re-adapts to the dangerous adult environment after spending their juvenile phase in a protected environment. The adult environment has an optimal policy and optimal value function. As the individual re-learns, its policy and estimated value function become closer to the optimal quantities. When these quantities of the individual match the optimal (within a degree of tolerance), then we say that the learning has been completed. The time, in number of steps from the start of adulthood to the completion of learning, is termed the re-learning time.

As the time spent in the juvenile phase increases, the time required to learn in the adult phase decreases (see Fig. 5).

**Fig. 5.**
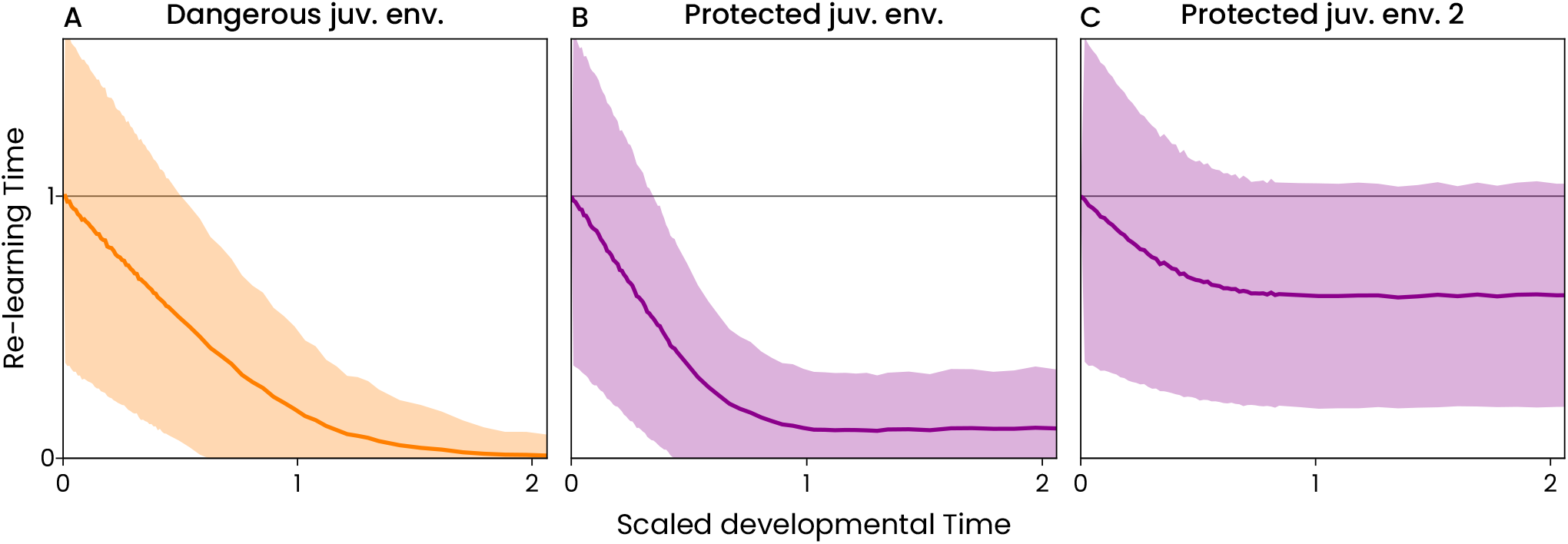
Dependence of relearning time in adulthood on the developmental time. **A**. The juvenile environment is as dangerous as the adult environment. (*ρ* = 0.9, *ϕ* = 0.6, Δ*E*_*inj*_ = 4) As developmental time increases, the time required for re-learning decreases. The line is almost linear, but deviates due to the stochastic nature of the process. **B**. The adult environment is the same as that in A, but the juvenile environment is protected with no injury cost 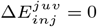. As the individual spends more time in the protected juvenile environment, the time for re-learning in the dangerous adult environment decreases. For this case, the re-learning time decreases faster than in the case of the dangerous environment. **C**. The adult environment is different from the left panels (*ρ* = 0.8, *ϕ* = 0.2, Δ*E*_*inj*_ = 4). The juvenile environment is protected compared to this, with no injury cost 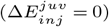. In this case, as the developmental time increases, the re-learning time does not continue to decrease. It stops improving at a very high re-learning time.

If the environment in the juvenile phase is the same as the adult, then the decrease in re-learning time with an increase in developmental time is nearly linear, with a deviation at high developmental times due to stochasticity. If, instead, the juvenile environment is different from the adult environment, then this curve is, in general, different. For some parameter values, the protected juvenile environment may accelerate relearning and make this curve fall more rapidly than before (Fig. 5B). For other parameter values, the re-learning time from a protected juvenile environment may still be very high even for long developmental times (Fig. 5C).

This curve can be approximated by an exponential decay *T*_*learn*_ ~exp(− *vT*_*dev*_). The decay parameter *v* can be considered as the learning speed. The higher this value is, the faster re-learning is in adulthood for low values of developmental time. Taking the ratio of the learning speed for a protected juvenile environment and that for a dangerous juvenile environment, we obtain the relative learning speed. We plot this value for a range of parameter values and protection level of the juvenile environment in Fig. 6. For high capture probability and high abundance of dangerous prey, the learning speed in a protected juvenile environment is higher than that in a dangerous environment. For low capture probability, the learning speed in a protected juvenile environment is, in general, lower than that in a dangerous juvenile environment. Even for low capture probability, there is a small region in parameter space with a high relative learning speed. These differences are only exaggerated as the juvenile environment becomes more protected.

**Fig. 6.**
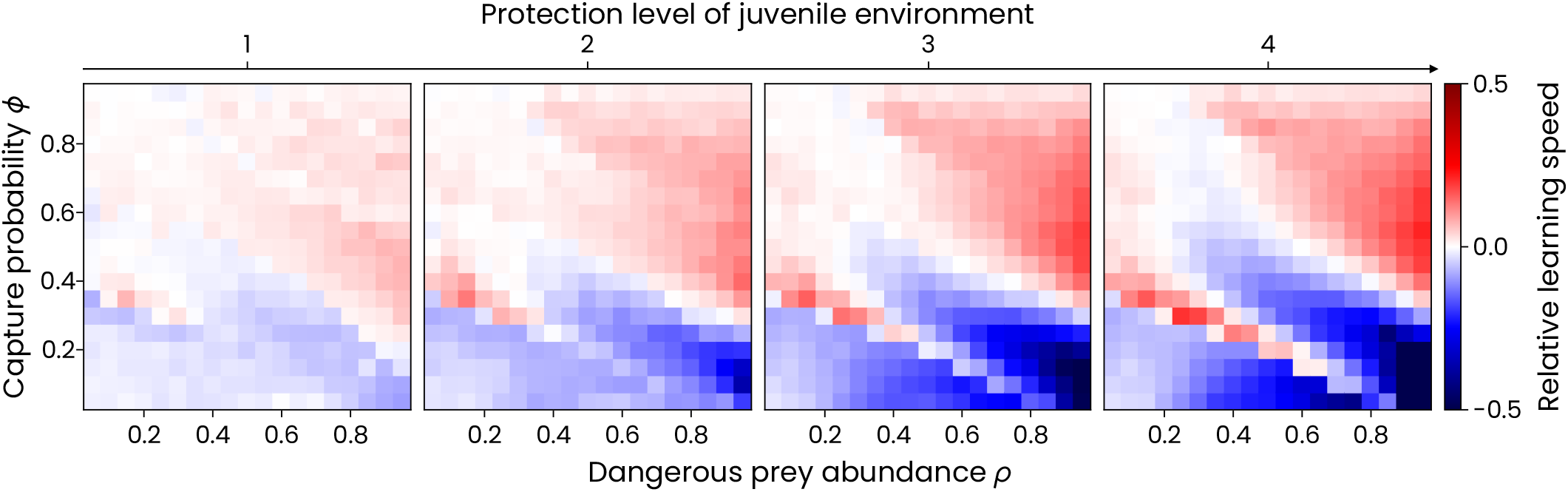
Relative learning speed across juvenile environment protection levels. Relearning time as a function of developmental time can be approximated and fitted with an exponential decay for developmental times less than 1. The ratio of the decay exponents of a protected and dangerous juvenile environment gives the relative learning speed in the protected environment. Relative learning speed is shown as a function of dangerous prey abundance (*ρ*), capture probability (*ϕ*) and juvenile environment protection level, measured by the reduction in injury cost from the dangerous to the protected environment 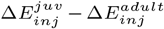. The adult phase has an injury cost of 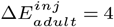. Positive values (red) indicate that a juvenile phase in a protected environment needs a shorter relearning time compared to a juvenile phase in a dangerous environment of the same developmental length. Negative values (blue) indicate that a protected juvenile phase results in a longer relearning time than an unprotected juvenile phase of a similar duration. The red and blue regions are roughly the same size and remain so even as the juvenile environment becomes more protected. However, they do become more prominent as the protection level increases. That is, for some parameters, the learning is highly accelerated, while for others, it becomes highly decelerated.

The relative learning speed gives us mainly information about what happens for low developmental times. However, the curve of learning time may deviate from exponential decay at higher developmental times. Thus, we plot the learning time for a range of developmental times in Fig. A.3. Here, a positive value means that the protected juvenile environment has a higher learning time than the dangerous juvenile environment, i.e. learning is slower in the protected juvenile environment compared to the dangerous juvenile environment. We see that the difference between protected and dangerous juvenile environments becomes higher as developmental time increases and as the juvenile environment becomes more protected. There is a large region in the parameter space at all levels of protection, where the learning time is still very high, even for long developmental times.

### 3.4. Benefit of protected juvenile environment

In this section we look at the overall benefit of a protected juvenile environment on learning. Learning can be benefited in the form of high relative adult performance or high relative learning speed. The relative adult performance can be high after a short juvenile phase or after an extended juvenile stage. Depending on the parameters of the environment and the protection level of the juvenile phase, the kind and level of benefit are different. This is shown in Fig. 7. We say that a protected juvenile environment provides one of these benefits, and the value for that metric for a protected juvenile environment is higher than that for a dangerous juvenile environment. For the metric of adult performance after an extended juvenile phase, we consider a benefit even if the protected environment matches the metric of the dangerous environment.

**Fig. 7.**
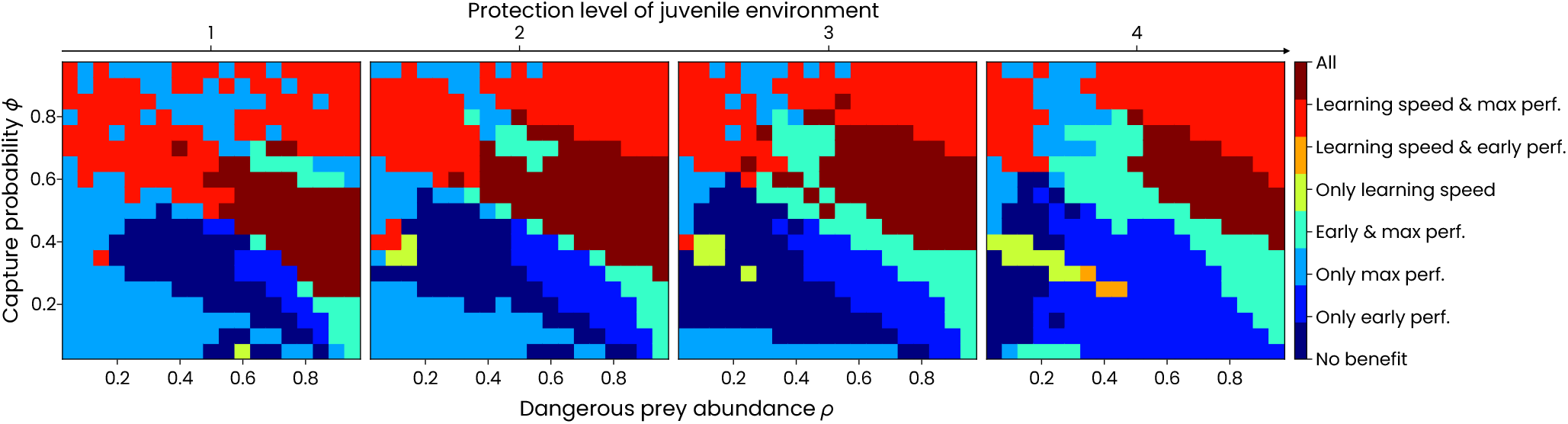
Benefit of protected juvenile environment. The benefit of a protected juvenile environment can either be in terms of a high (a) adult performance after a short juvenile phase (early perf.), (b) adult performance after an extended juvenile phase (max perf.), or (c) learning speed, or a combination of these. The type and extent of benefit changes across the environmental conditions of dangerous prey abundance (*ρ*), capture probability (*ϕ*) and protection level of juvenile environment (quantified by reduction in cost of injury).

For a single environmental condition, there can be one or more of these benefits. As the parameters of the environment and of the protection change, the kind and extent of the benefit change in highly non-linear ways. This is because these benefits depend on the optimal policies and values of the dangerous and protected environments, their properties and their extent of match or mismatch. Since these quantities vary in highly non-linear ways, so do the metrics of learning benefits. However, there are some general patterns. The region with no learning benefit first increases in size and then decreases as the protection level of the juvenile environment is increased.

For a highly protected environment, this region is very small. That is, there is some kind of benefit to learning for all environmental conditions. There is a sizeable region where only the adult performance for an extended juvenile phase is benefited. This region shrinks in size as the protection level increases. The regions of no benefit and only maximum performance benefit are taken over by the region with only an early performance benefit. For higher protection levels, a small region with a benefit in both early and maximum adult performance becomes prominent for a slightly higher capture probability than the previously mentioned regions. For high capture probability, there are two regions, one in which learning is benefited in the form of a high learning speed and high maximum performance, and the other in which learning is benefited in all manners. These regions are roughly the same size for all protection levels.

## 4. Discussion & Conclusion

We have investigated the effect of juvenile play on learning and adaptation towards a dangerous environment, and on performing risky behaviour. A parent can provide their juvenile offspring a protected environment, where they can learn and adapt to the dangerous environment they will face in adulthood. We have quantified the effect of a protected juvenile environment on its learning and adaptation through multiple metrics for different environmental conditions and protection levels of the juvenile environment. Our work shows that depending on the environmental conditions and the protection level of the juvenile environment, the kind and extent of benefit on learning vary substantially. The relation between the environmental conditions is highly non-linear. Theory from the field of transfer learning tells us that these metrics of learning depend on the optimal value function *V* *(*s*), the optimal policy *π*\*(*s*), the transition function *P* (*·*|*s, a*) of the two different environments and how they are related to each such, such as their distance (44). However, we can only analytically derive bounds for these quantities and not explicit relations.

To ensure the development of well-adapted adults, it is not sufficient to just provide a protected juvenile environment. This can be understood by analysing the ways in which a protected environment can benefit learning. One of the ways in which learning is benefited is with a high adult performance after an extended juvenile phase. This means that if the juvenile learns for a sufficiently long time in the protected juvenile phase, then it will perform close to optimum as an adult in the real dangerous environment. Such a high maximum adult performance is seen in a large region of the parameter space of the environment. When the abundance of dangerous prey and capture probability are high, a protected environment allows the juvenile to adapt to the dangerous environment. For these conditions, the simple parenting strategy of providing a completely protected environment for the entire duration of the juvenile phase will produce well-adapted adults. For other conditions, such a simple parenting strategy is not sufficient and will result in incompletely adapted juveniles being exposed to the dangerous adult environment.

Another form of benefit is a high adult performance after a short juvenile phase. If the adult performance after a short protected juvenile phase is higher than that of the dangerous juvenile phase, it means that a protected environment results in rapid adaptation early in the juvenile phase. The final form of benefit to learning that we study is a high learning speed. Switching from a protected to a dangerous environment requires some level of re-learning and re-adaptation. Having a high relative learning speed means that for the same developmental time, the re-learning time is reduced. There are sizeable regions of the parameter space where a protected juvenile environment provides either or both of these benefits. Having such benefits still might allow for proper adaptation, but requires a more involved parenting strategy. Providing a highly protected environment initially can result in rapid initial adaptation due to the high early adult performance and high learning speed. But this cannot provide complete adaptation. Gradually decreasing the protection level until the juvenile environment matches the adult environment can result in better adaptation due to a better match between the juvenile and adult environments. This is seen in meerkats, which hunt deadly scorpions, where mothers bring increasingly dangerous scorpion prey to their offspring as they grow and develop (14). Thus, more complex and environment-dependent parental care may be required in order to produce individuals adapted for adulthood. There may even exist an optimal parental care strategy which results in rapid adaptation with a low juvenile mortality. Further investigation is required in this aspect.

We have studied the case where individuals learn by themselves through experience. Several species that display high levels of learning also display high levels of social and group behaviours (45–47). In such species, individuals have another strategy to utilise for adaptation — social information (48, 49). Naive juveniles can learn from the experience of their peers. However, the best social learning strategy in such situations is not well known, and this forms another aspect of future study.

The learning process itself can evolve and increase in complexity in order to account for such a difference in juvenile and adult environments. The information about both environments can be used in order to derive a relation between them and convert the knowledge and experience acquired in a protected environment to apply in a dangerous environment. This can result in rapid re-adaptation after learning in a highly protected juvenile environment, even with a mismatch between both environments. Such a process can be further studied using the approaches from the field of transfer learning (50).

Here, we assume that learning happens uniformly across the lifetime of the individual. However, it is costly to maintain the cognitive mechanisms required for learning throughout the lifetime. Animals would therefore evolve to be more sensitive to stimuli and would learn more within the early periods of their (51). We use a learning algorithm in which the action to be taken for each energy level is stored separately. A high level of cognitive capability required by such a learning mechanism would come with high costs. It is possible that these high costs would lead to the evolution of a mechanism with less precision and more generalised rules (52). However, regardless of the coarse-grained nature of the algorithm, the results observed from our model still apply. The rewards obtained by the animals are not obtained from the environment, but rather through an internal mechanism that provides reward signals (37). Therefore, evolution needs to generate an appropriate reward system which has the property that higher reward signals in general leads to a higher expected fitness. We created and analysed this model with the assumption that there exists such an internal reward system capable of encouraging the optimal behaviour.

There are several complications which we have simplified in the model for the sake of analysis. The abilities of animals would also change over the course of their life, and ignoring this was another simplifying assumption we made. The decision process of an animal is also much more complex than depicted here. There are several steps and decisions taken by the predator to go from spotting prey to a full-on attack. A predator may take note of its size and energy level as it approaches, whether it is in a group or not. A predator may also abandon a prey mid-attack if it senses too much risk. Several predators also utilise innovative attacks to deal with the dangerousness of prey.

A protected environment can allow naive juveniles to adapt to a dangerous environment without a high juvenile mortality. However, more complex parental care strategies are required to make this possible under all environmental conditions and to make the learning optimal. Many further investigations are required in order to understand this phenomenon better.

## Code and Data Availability

All simulation code, datasets and plotting scripts are available on Zenodo at https://doi.org/10.5281/zenodo.15857802.

## ACKNOWLEDGEMENTS

The authors thank members of the `T-Eco-Evo` group for inspiring discussions. Funding support from Julius-Maximilians-University Würzburg is gratefully acknowledged.

## Appendices

### A. Model details

#### A.1. Decision process

The jump process (*E*_*n*_, *t*_*n*_), where Δ*t* is the time between the transitions and *n* is the jump number, is a semi-Markov decision process (SMDP) because of the following features. The next state and jump (or transition) time only depend on the preceding state and action of the predator. The jump time is not a constant but can continuously vary. It is only semi-Markov because the jump times do not follow an exponential distribution as required by a fully Markov continuous time Markov decision process (CTMDP).

Table A.1 shows the possible outcomes and their probabilities for the three compound behaviours.

**Table A1.** Outcomes and their probabilities for the different compound behaviours the predator chooses.

| Behaviour | Outcome / Probability |  |  |  |
| --- | --- | --- | --- | --- |
| | Capture safe<br>$+\Delta E_{safe}$<br>$\Delta t_{search} + \Delta t_{safe}$ | Capture dang.<br>$+\Delta E_{dang}$<br>$\Delta t_{search} + \Delta t_{dang}$ | Incur injury<br>$-\Delta E_{inj}$<br>$\Delta t_{search} + \Delta t_{inj}$ | No attack<br>$-m$<br>$\Delta t_{search}$ |
| Indiscriminate | $(1 - \rho)s_{safe}$ | $\rho\phi s_{dang}$ | $\rho(1 - \phi)s_{inj}$ | 0 |
| Risk-prone | 0 | $\rho\phi s_{dang}$ | $\rho(1 - \phi)s_{inj}$ | $(1 - \rho)s_{skip}$ |
| Risk-averse | $(1 - \rho)s_{safe}$ | 0 | 0 | $(1 - \rho)s_{skip}$ |

For our analysis of the decision process, we consider a stochastic policy. That is, *π*(*A|E*) is the probability of taking action *A* at energy level *E*. For a given policy *π*, the value function can be calculated with the recursive set of equations:

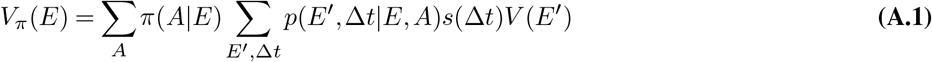

where, *p* is the transition probability function, *s* is the survival probability. The state *E* = *E*_*c*_ is the only state with any reward and always gets a constant reward. Thus, we can take the value of this state to be *V* (*E*_*c*_) = 1 and calculate the value of the other states relative to this. The optimal value is *V* * = sup_*π*_ *V*_*π*_, and the optimal policy is *π*\* such that *V*_*π*_*\** = *V* *. The optimal value function is found using the Bellman optimality equations:

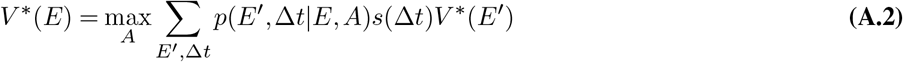

The optimal policy, the policy with the highest fitness, can be calculated from this optimal value as:

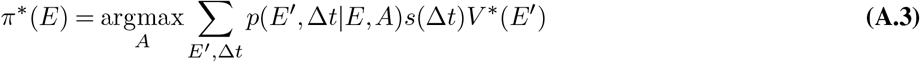

Therefore, we have a variable discount factor *s*(Δ*t*), which depends on the transition time between states.

##### Choosing of parameters

The parameters of the decision process were chosen such that a wide range of behaviours and optimal policies can be observed by varying just the two parameters of dangerous prey abundance *ρ* and capture probability *ϕ*. We set *E*_*c*_ = 20 to ensure that learning converges to the optimal policy within a reasonable computational time while displaying a wide range of optimal policies. The energy parameters Δ*E*_*i*_ need a sufficiently high variance to observe a realistic optimal policy without rapid switching between actions for neighbouring energy levels. This is why we set the energy parameters as random normal variables left-truncated at zero. The parameters of the decision process are listed in Table A.2.

**Table A2.** All the quantities and their values as used in the model. The energy quantities Δ*E*_*i*_ are random normal variables 𝒩 (*µ*_*i*_, 1.5*µ*_*i*_), which are left-truncated at 0, with *µ*_*i*_ shown below.

| Symbol | Meaning | Value(s) |
| --- | --- | --- |
| $\rho$ | Dangerous prey abundance | 0-1 |
| $\phi$ | Dangerous prey capture probability | 0-1 |
| $\Delta E_{inj}^{adult}$ | Energy cost of injury in adulthood | 4 |
| $\Delta E_{inj}^{juv}$ | Energy cost of injury in juvenile phase | 0, 1, 2, 3, 4 |
| $E_c$ | Maximum energy level | 20 |
| $\Delta E_{safe}$ | Energy benefit of safe prey | 2 |
| $\Delta E_{dang}$ | Energy benefit of dangerous prey | 10 |
| $m$ | Metabolic energy cost | 1 |
| $\Delta t_{search}$ | Searching time | 1 |
| $\Delta t_{safe}$ | Handling time of safe prey | 1 |
| $\Delta t_{dangerous}$ | Handling time of dangerous prey | 1 |
| $\Delta t_{inj}$ | Recovery time after injury | 1 |
| $\alpha$ | Learning rate | 0.01 |
| $\epsilon$ | Exploration rate | 0.2 |
| $d$ | Background mortality rate | 0.1 |
|  | Learning convergence threshold | 0.25 |
|  | Number of learning experiments | 10 000 |
| | Dynamic programming convergence threshold | $10^{-6}$ |

### A.2 Dynamic Programming

Dynamic programming (26) is an iterative algorithm to numerically solve the equations A.2, A.3 and obtain the optimal value and policy. We use the value iteration algorithm to estimate the optimal value *V* *, which can be summarized as follows. For each *E* = 0, 1,…, *E*_*c*_ calculate the following and update the estimate of *V* *(*E*)

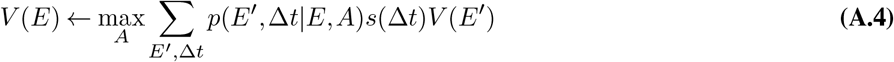

This calculation is repeated until the changes to the value function in each update become less than a desired threshold, which we set to be 10*−*6. With the calculated estimate for *V* *(*E*), we use Eq. A.3 to obtain the optimal policy.

### A.3. Reinforcement Learning

We use temporal-difference reinforcement learning to model learning. In this form of learning, we use an action-value function *Q*_*π*_(*E, A*). It is the expected total reward received on taking action *A* in state *E* following policy *π* thereafter. It is related to the value function *V* as follows

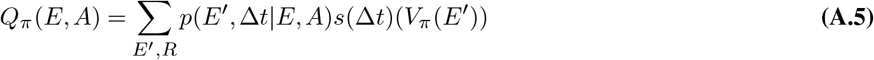

Here, *V*_*π*_(*E*_*c*_) = 1, as a reward is obtained only upon reaching the highest energy level. Since the total number of states and actions are discrete and finite, we use a table to keep track of the estimate of *Q*(*E, A*) for each state *E* and action *A*.

The predator updates its estimates of the optimal value as it experiences the environment, its state transitions and rewards. Suppose the agent has energy level *E* and takes an action *A*, resulting in a transition to the energy level *E*^*′*^ after a time Δ*t*. The agent then follows its policy and takes the next action *A*^*′*^. Then, the parameters are updated as follows:

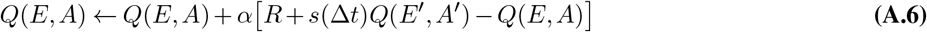

Here, we replace the otherwise constant discount factor with *s*(Δ*t*) = exp(− *d*Δ*t*) (53). By doing this, we assume that the individual’s learning process has some knowledge about the background mortality in the environment and knows how to discount future rewards. This information could be gained evolutionarily.

The agent follows an *ϵ*-greedy policy, allowing it to be exploitative and explorative. It is mathematically defined as follows. For every energy level *E*, let *A*\* = argmax_*A*_ *Q*(*E, A*), and *n*_*A*_ be the number of actions. The *ϵ*-greedy policy is

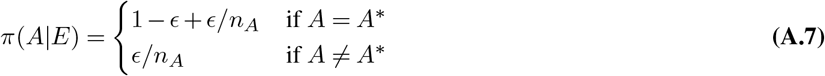

#### Choosing of hyperparameters

The learning hyperparameters — the learning rate *α* and exploration rate *ϵ* — need to be carefully chosen to observe convergence to the optimal solution and show consistent and sufficiently low learning times. *α* should be sufficiently low to ensure convergence (close) to the optimal policy. If it is too high, the policy or value can have large jumps between steps, causing it to reach the optimal very slowly and inconsistently, if at all. *ϵ* should be low, but not too low, to promote sufficient exploration and prevent the policy from getting stuck at a local optimum. It should also not be too high, otherwise the agent continues to explore even when it is close to the optimum, which reduces its performance. We choose *α* = 0.01, and *ϵ* = 0.2, as they provide convergence to the optimal policy with consistent learning times. These learning times are not very large, so all the required simulation runs can be performed reasonably.

For calculations of learning time, we compare the learnt value and policy functions against the optimal value and policy functions, and stop the learning process and say that learning has been completed when the error value drops below a certain threshold (we chose 0.25). The error is calculated as follows:

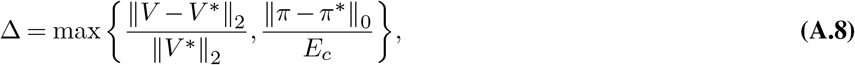

where, *V* and *π* are the current estimated value and policy, while *V* * and *π*\* are the optimal value and policy. ∥·∥_2_ represents the Euclidean distance, and ∥·∥_0_ represents the Hamming distance.

Figure. A.1A shows that the learning agent reaches within this error threshold of the optimal policy and value. The time taken to reach the optimal policy and value starting from a random policy depends on the parameters of the environment, as shown in Fig. A.1B

**Fig. A1.**
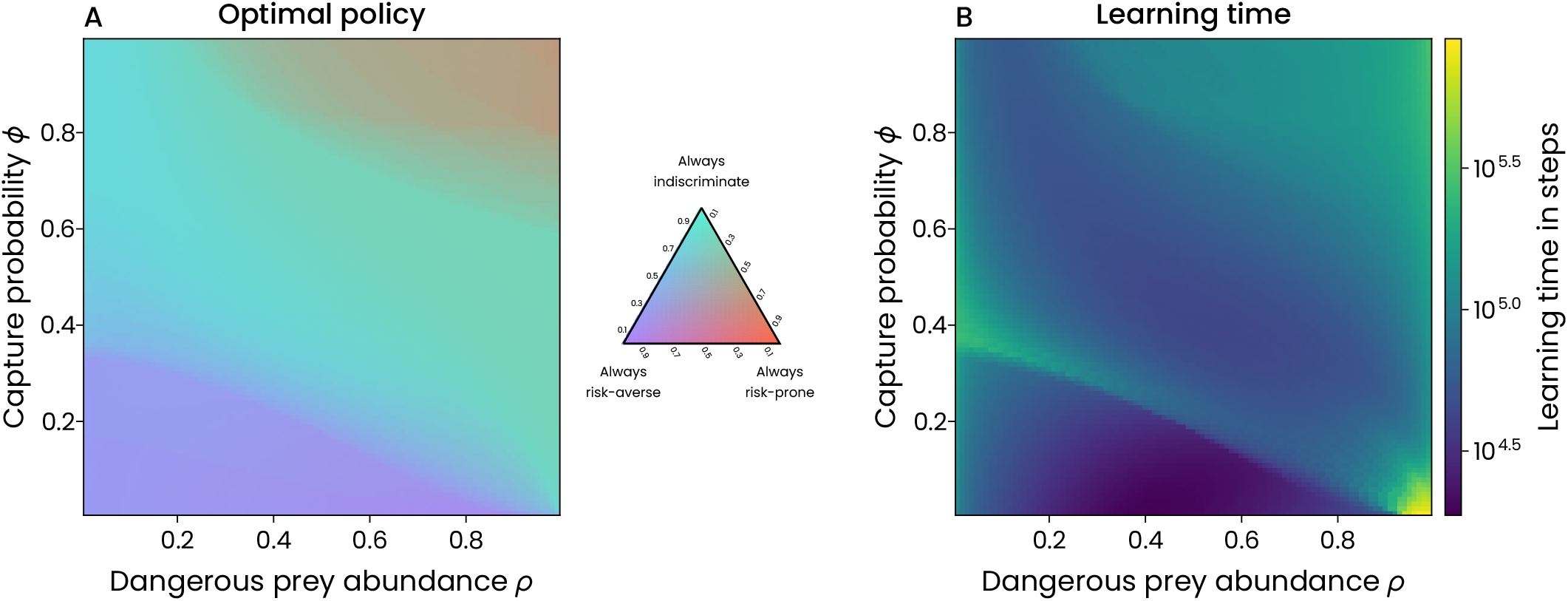
Optimality of reinforcement learning. **A**. The policy reached by a learning agent after gaining sufficient experience through learning, for different dangerous prey availability *ρ* and capture probability *ϕ*. **B**. The learning time in steps required for the learning agent to converge to the policy shown in A. This shows that a reinforcement learning agent can reach the optimal policy with the chosen hyperparameters of learning, and that the number of steps to reach optimality varies between environmental conditions.

### A.4. Results

Figures A.2 and A.3 show the metrics of relative adult performance and relative re-learning time for various developmental times and protection levels of the juvenile environment. This shows how the different regions of high performance or low learning time change and move as the developmental time increases or the protection level increases.

**Fig. A2.**
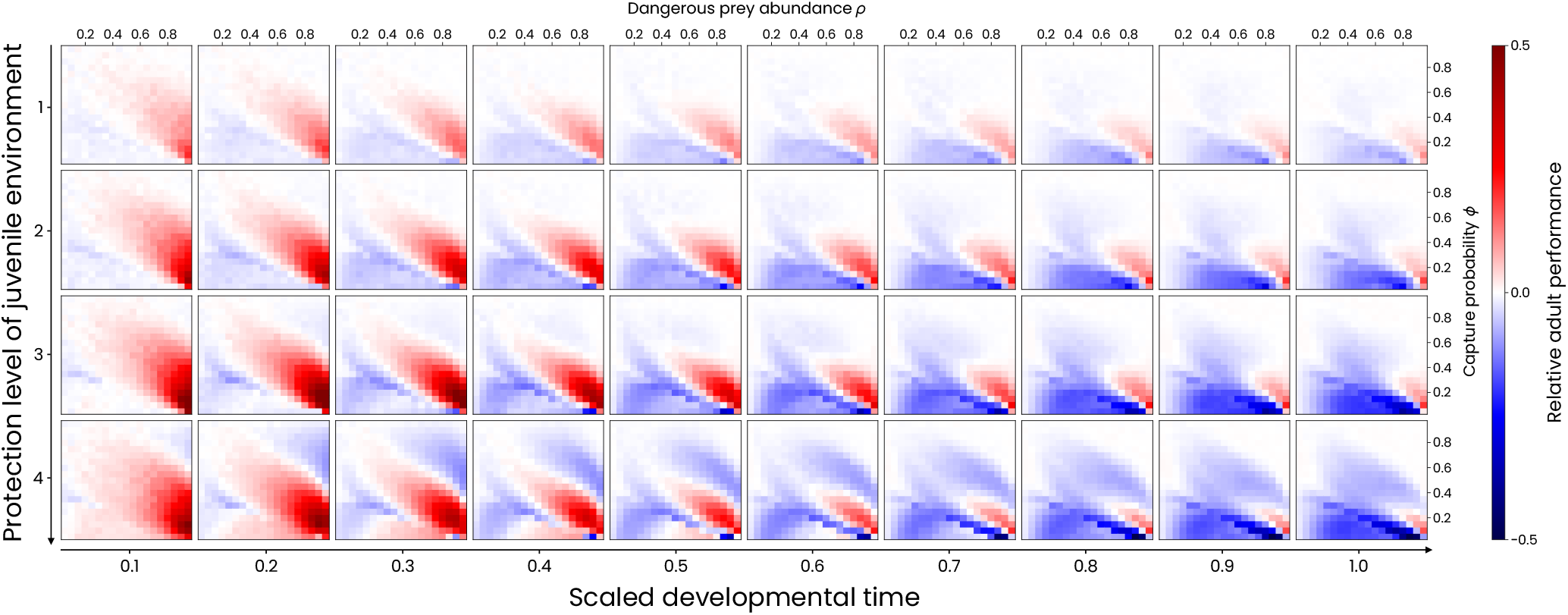
Relative adult performance across developmental times and juvenile environment protection level. We plot the performance in a protected environment relative to its corresponding dangerous environment for different developmental times, parameters of the environment (*ρ, ϕ*) and protection level of the juvenile environment. From left to right, each heatmap is for a different developmental time. From top to bottom, each heatmap is for a different level of protection in the juvenile environment. Within each heatmap, performance is plotted over a grid of dangerous prey abundance *ρ* and capture probability *ϕ*. Red regions indicate parameter combinations where the protected juvenile environment leads to higher adult performance; blue regions indicate the opposite. White regions denote comparable adult performance following both protected and dangerous juvenile environments. Protective environments often yield better adult performance across a broad parameter space for shorter developmental times. As developmental time increases, this advantage diminishes, with blue regions growing and red regions shrinking. The relative benefit of a protected juvenile phase depends strongly on both environmental parameters and the protection level of the juvenile environment.

**Fig. A3.**
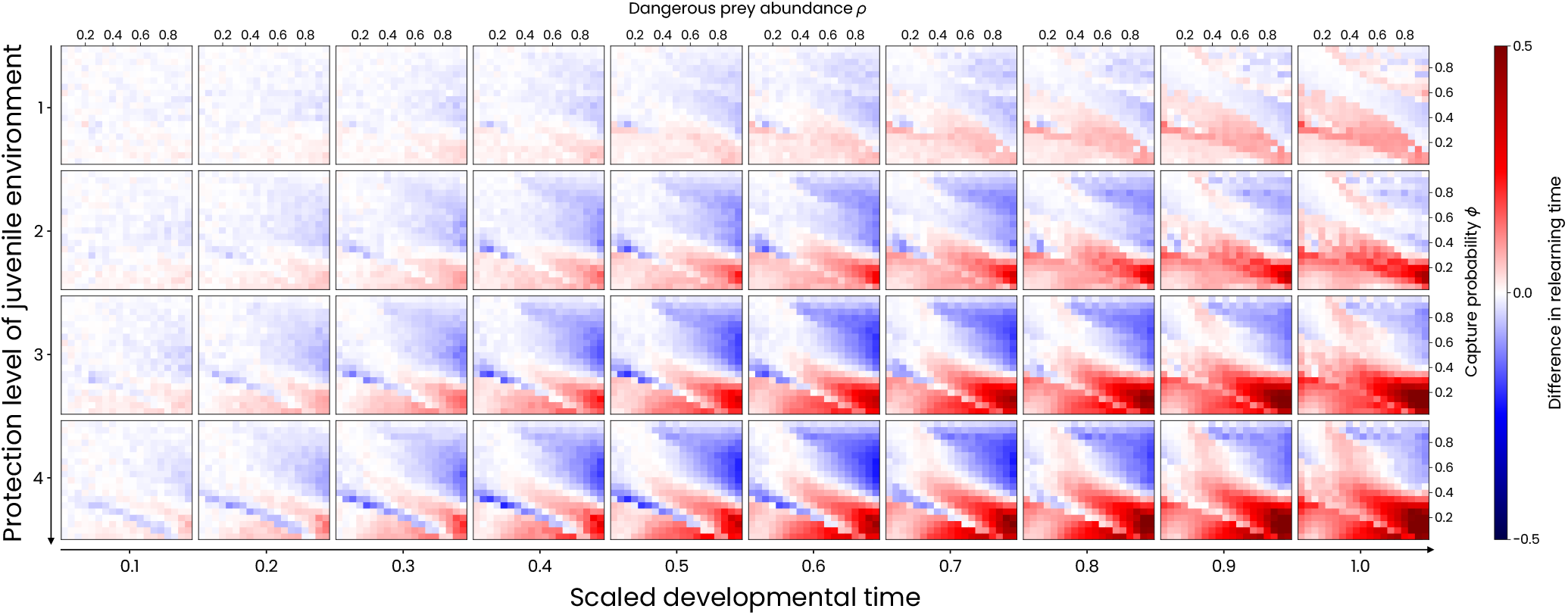
Relative learning time across developmental times and juvenile environment protection levels. We plot the re-learning time after a protected juvenile phase relative to its corresponding dangerous environment for different developmental times, environmental parameters (*ρ, ϕ*) and protection level of juvenile environment 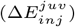. In two regions, re-learning time in a protected environment is either lower than (blue) or higher than (red) that of a dangerous environment. These regions become more prominent and larger for higher developmental times and more protected juvenile environments. For high developmental times, the region with higher relearning time becomes more prominent and larger than the other region.

